# Descending somatosensory and motor cortical inputs shape auditory processing in the midbrain

**DOI:** 10.64898/2026.08.25.747140

**Authors:** Hae Yeon Kang, Jisoo Han, Miguel Sánchez-Valpuesta, Joonyeol Lee, Seong-Gi Kim, Gunsoo Kim

## Abstract

Integrating multisensory and behavioral information is essential for sensory perception. In the auditory system, multisensory and behavioral influences emerge early in subcortical structures. Descending projections from non-auditory cortical areas are well positioned to convey such signals, yet how they shape subcortical auditory processing remains poorly understood. Here, we investigated corticocollicular projections from the primary somatosensory (S1) and motor (M1) cortices to the inferior colliculus (IC), a principal integration center in the auditory midbrain. We found that trunk- and limb-related regions of S1 and M1 form prominent monosynaptic projections to the IC, and that optogenetic activation of these projections robustly drives IC activity. Notably, a substantial population of cortical-responsive neurons did not respond to sound. In sound-responsive neurons, concurrent cortical stimulation enhanced sound-evoked responses, whereas cortical activation preceding sound onset suppressed them. Furthermore, both cortical-responsive IC neurons and deep-layer S1 and M1 neurons exhibited locomotion-related modulation and anticipatory activity prior to movement onset, suggesting that these descending pathways convey movement-related signals to the IC. Together, our findings identify a descending sensorimotor circuit that integrates body- and movement-related information with auditory processing in the auditory midbrain.

## Introduction

Integrating multisensory and behavioral information is essential for sensory perception. In the auditory system, neural representations are shaped not only by acoustic input but also by signals from other sensory modalities and behavioral state^1–5^. Multisensory and behavioral influences on auditory processing have been well characterized in the auditory cortex^6–18^. However, subcortical auditory structures also exhibit multisensory^19–22^ and movement-related activity^23–30^. This subcortical integration likely arises in part from ascending crossmodal inputs, but also depends on descending projections from multiple cortical areas^31–34^. Yet how these descending non-auditory inputs shape auditory processing in subcortical circuits remains poorly understood.

The inferior colliculus (IC) is the principal midbrain hub of auditory processing, where virtually all ascending auditory inputs first converge^35–37^. While the central nucleus of the IC (CNIC) is regarded as predominantly auditory, somatosensory responses are especially prominent in the surrounding shell regions^21,22,38–40^. Along with ascending somatosensory inputs^41^, the IC receives extensive corticocollicular projections, providing an additional route through which non-auditory information can influence auditory processing. Prior studies have shown that, besides the auditory cortex^42–44^, corticocollicular projections also arise from multiple non-auditory cortical areas, including the primary somatosensory (S1) and motor (M1) cortices^31,32,34,45^. Whereas auditory corticocollicular projections are well known to shape sound-evoked responses^46–49^ and support sound-guided behavior^50,51^, comparatively little is known about how non-auditory corticocollicular projections influence auditory processing.

A recent study demonstrated that multiple non-auditory cortical regions, including S1 and M1, provide direct excitatory input to neurons in the CNIC^34^. Because those experiments measured only responses evoked by cortical stimulation, however, they did not address how these descending inputs influence auditory processing or contribute to sensorimotor integration in the IC. The strongest evidence that a non-auditory corticocollicular pathway shapes auditory processing comes from whisker-related S1 projections, which suppress sound-evoked responses in the auditory thalamus via inhibitory neurons in the IC shell^33^. While this demonstrates an important non-auditory cortical influence on auditory processing, it remains unknown whether descending sensorimotor corticocollicular pathways primarily modulate auditory responses or more broadly contribute to sensorimotor integration in the IC during behavior.

Here we investigated the organization and function of corticocollicular projections arising from S1 and M1. We show that trunk- and limb-related regions of S1 and M1 form prominent monosynaptic projections to the IC. Optogenetic activation of these projections robustly drove IC neurons and modulated auditory responses in a timing dependent manner. Furthermore, recordings in behaving mice suggest that these corticocollicular pathways convey movement-related signals to the IC during locomotion. Together, our findings identify a descending sensorimotor circuit that integrates body- and movement-related information with auditory processing in the auditory midbrain.

## Results

### Monosynaptic corticocollicular projections from trunk- and limb-related S1 and M1

We mapped corticocollicular projections by injecting a retrograde virus (AAV2-retro) expressing red fluorescent protein into the left IC (Fig. 1a). Three-dimensional reconstruction aligned to the Allen Mouse Brain Reference Atlas^52^ revealed dense labeling in the primary somatosensory (S1) and motor (M1) cortices, as well as several other non-auditory cortical regions, consistent with recent reports^32,34,45^ (Fig. 1b and Supplementary Fig. 1). In coronal sections, IC-projecting neurons were concentrated in cortical layer 5 and were located predominantly in trunk- and limb-related regions of S1 and M1 (Fig. 1c).

**Figure 1.**
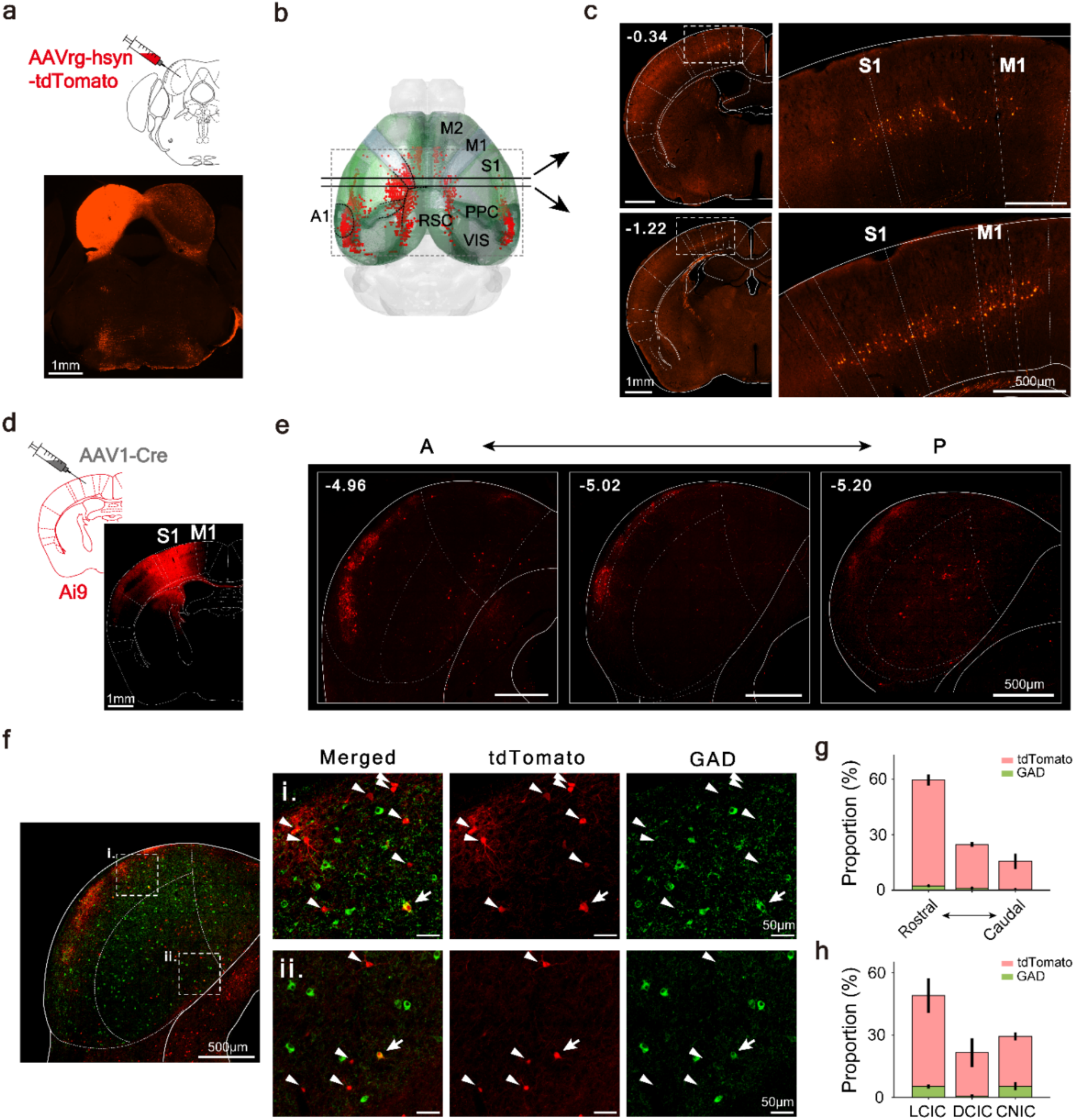
Monosynaptic corticocollicular projections from S1 and M1 to the IC. **a**) Schematic (top) and representative coronal section (bottom) showing the injection site of a retrograde AAV in the left IC. **b**) Three-dimensional reconstruction of retrogradely labeled cortical neurons (red dots) aligned to the Allen Brain Atlas. Horizontal lines indicate the rostrocaudal levels of the coronal sections shown in (**c**). Black dotted lines delineate cortical subregions; the auditory cortex (A1) is outlined with a dashed line. M2: secondary motor cortex; M1: primary motor cortex; S1: primary somatosensory cortex; PPC: posterior parietal cortex; VIS: visual cortex; RSC: retrosplenial cortex. **c**) Coronal sections showing IC-projecting neurons in S1 and M1. Right: higher magnification views of the boxed regions showing layer 5 corticocollicular neurons. **d**) Schematic of the anterograde transsynaptic tracing strategy. AAV1-Cre was injected into S1 of Ai9 reporter mice to label recipient IC neurons. **e**) Representative coronal sections at 3 rostrocaudal levels, showing transsynaptically labeled recipient neurons and cortical axons in the IC. **f**) GAD67 immunolabeling of S1-recipient IC neurons. Left: low-magnification confocal image showing tdTomato+ recipient neurons (red) and GAD67+ neurons (green). Right: higher-magnification images of the boxed regions (i, ii), showing merged and individual fluorescence channels. Arrows indicate GAD67+ recipient neurons; arrowheads indicate GAD67-recipient neurons. **g,h**) Distribution of recipient neurons along the rostrocaudal axis of the IC (**g**) and across IC subdivisions (**h**). Recipient neurons labeled from S1 and M1 injections were pooled for quantification. Red: GAD67-neurons; green: GAD67+ neurons. DCIC: dorsal cortex of the IC; LCIC: lateral cortex of the IC; CNIC: central nucleus of the IC. Data are presented as mean ± SEM (n = 4 mice).

To identify IC neurons receiving these cortical inputs, we injected an anterograde transsynaptic virus (AAV1) expressing Cre recombinase into S1 or M1 of Ai9 reporter mice^53^ (Fig. 1d). Labeled recipient neurons were distributed throughout all major IC subdivisions, with the highest density in the lateral cortex (LCIC), but were also present in the central nucleus (CNIC) and dorsal cortex (DCIC) (Fig 1e,h and Supplementary Fig. 2). Quantification across 3 coronal levels revealed a rostrocaudal gradient, with recipient neurons most abundant in the rostral IC (Fig. 1g).

Within the LCIC, recipient neurons were observed both within and outside patch-like regions enriched for glutamic acid decarboxylase^31^ (GAD67). Co-labeling revealed that the vast majority of recipient neurons were GAD67-negative, with only ∼5% expressing GAD67 (n = 4 mice). The small population of GAD67-positive recipient neurons was largely confined to the rostral LCIC and CNIC (Fig. 1f-h).

Together, these viral tracing results demonstrate that trunk- and limb-related S1 and M1 establish direct monosynaptic connections with neurons throughout the IC. The predominance of GAD67-negative recipient neurons further suggests that the descending sensorimotor cortical inputs primarily target excitatory IC neurons.

### Optogenetic activation of IC-projecting S1 and M1 neurons drives IC activity

In contrast to descending inputs from the auditory cortex, the functional properties of non-auditory cortical inputs to the IC have not been extensively investigated^34,47,54–56^. To selectively activate IC-projecting cortical neurons, we injected a retrograde AAV encoding channelrhodopsin-2 (ChR2) into the left IC (Fig. 2a).

**Figure 2.**
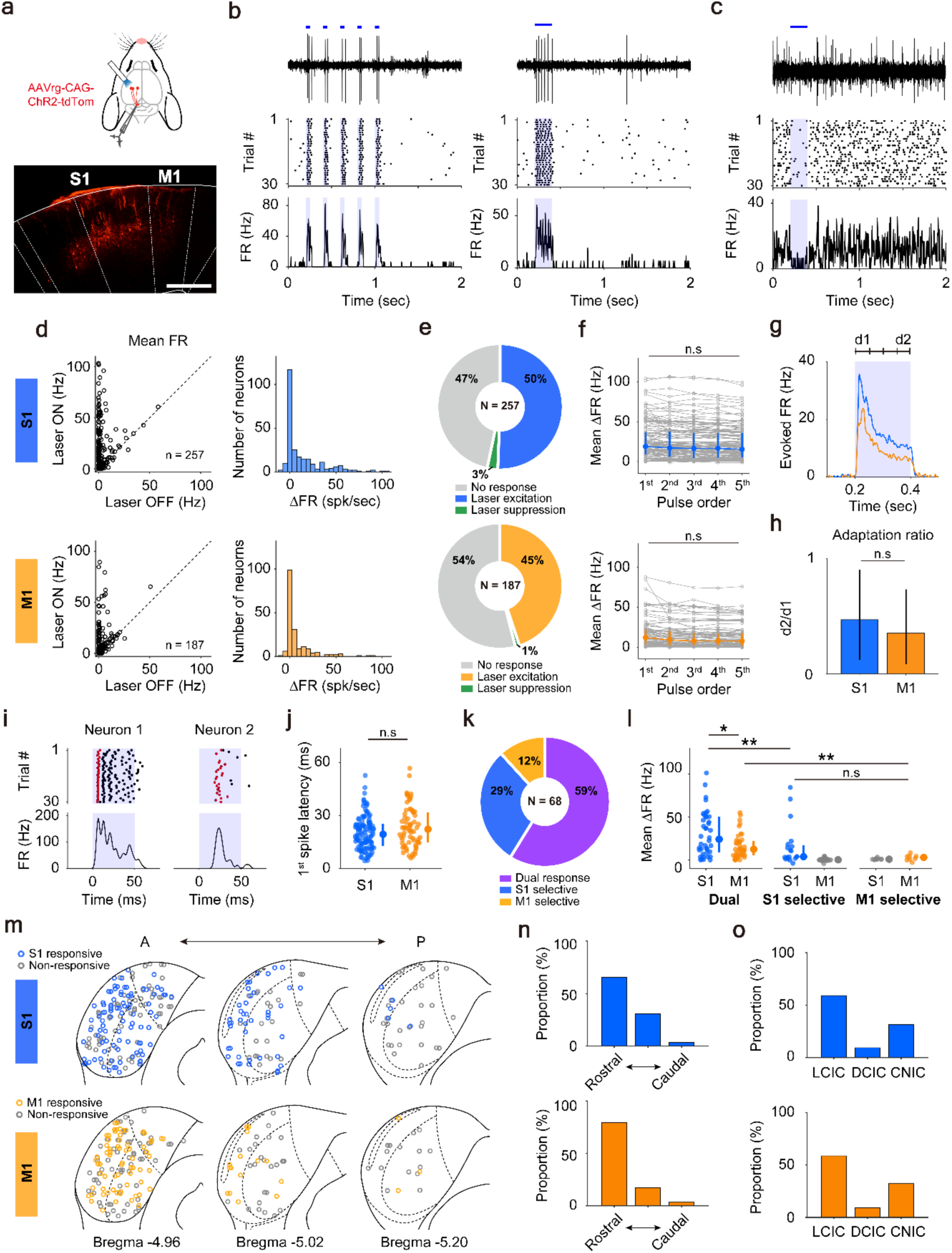
Optogenetic activation of IC-projecting S1 and M1 neurons drives IC activity. **a**) Schematic of the experimental approach (top). IC-projecting cortical neurons, which express ChR2-tdTomato, were optogenetically stimulated while recording neural activity in the IC. Bottom: Representative fluorescent image showing ChR2-tdTomato expression in S1 and M1. **b**) Example IC neuron excited by optogenetic stimulation of S1 with a 5 Hz pulse train (five 50 ms pulses, left) or a single 200 ms pulse (right). Spike waveform (top), raster plot (middle), and peri-stimulus time histogram (PSTH; bottom) are shown. Blue horizontal lines and shading indicate periods of laser stimulation. **c**) Example IC neuron suppressed by S1 stimulation. **d**) Scatter plots comparing firing rates between laser-OFF and laser-ON conditions (left) and distribution of firing rate changes (ΔFR, right) across IC neurons during S1 (top) and M1 (bottom) stimulation. Firing rates were averaged across the 5 pulses of 5 Hz stimulation train (S1: p = 1.93 x 10^-23^, n = 257; M1: p = 1.53 x 10^-14^, n = 187, Wilcoxon signed-rank test). FR: firing rate. **e**) Proportion of IC neurons showing excitation, suppression, or no response to S1 (top) or M1 (bottom) stimulation. **f**) IC responses during the 5 Hz pulse train. Mean firing rate changes (ΔFR) did not differ across the 5 pulses (S1: p = 0.63, n = 134; M1: p = 0.33, n = 84, Kruskal-Wallis test). **g**) Population PSTHs of responses to a 200 ms stimulation of S1 (blue) or M1 (orange). Shading indicates the stimulation period; d1 and d2 indicate the initial (0-50 ms) and late (150-200 ms) response epochs, respectively. **h**) Adaptation ratio, calculated as the ratio of mean firing rates during the late and initial epochs (d2/d1), did not differ between S1 and M1 stimulation (S1: median = 0.47; M1: median = 0.35; p = 0.33, Wilcoxon rank-sum test). **i**) Example IC response to S1 stimulation, where first spike latency was measured. Red dots indicate the first spike on each trial, and blue shading indicates the 50-ms laser stimulation period. **j**) Comparison of first-spike latencies evoked by S1 and M1 stimulation (median latency: S1: 19.4 ms, M1: 22.3 ms, p = 0.09, Wilcoxon rank-sum test). **k**) Proportion of S1-selective, M1-selective, and dual-responsive IC neurons. **l**) Mean responses evoked by S1 and M1 stimulation in dual responsive, S1-selective, and M1-selective IC neurons (p = 0.021, S1 vs. M1 responses in dual-responsive neurons; p = 0.009, S1 responses in dual-responsive vs. S1-selective neurons; p = 0.0019, M1 responses in dual-responsive vs. M1-selective neurons; p = 0.27, S1 responses in S1-selectrive vs. M1 responses in M1-selective neurons, Wilcoxon rank-sum tests). **m**) Spatial distribution of S1-responsive (top, blue) and M1-responsive (bottom, orange) IC neurons across 3 rostrocaudal levels. Gray circles indicate non-responsive neurons. **n**) Proportion of S1-responsive (top) and M1-responsive (bottom) neurons along the rostrocaudal axis of the IC (S1: n = 137 neurons; M1: n = 87 neurons). **o**) Proportion of S1-responsive (top) and M1-responsive (bottom) neurons across IC subdivisions.

In anesthetized mice, optogenetic stimulation of IC-projecting neurons in the hindlimb and forelimb regions of the left S1 reliably evoked responses in ipsilateral IC neurons (Fig. 2b-d; Supplementary Fig. 3a-d). Approximately half of the recorded IC neurons responded to S1 stimulation, with 50% showing excitation (n = 130 of 257 neurons) and only 3% showing suppression (n = 7 of 257 neurons), resulting in a population response distribution skewed toward excitation (Fig. 2d,e; Shapiro-Wilk test, p < 0.001). Similarly, stimulation of IC-projecting M1 neurons evoked excitatory responses in 45% of IC neurons (n = 84 of 187 neurons) and suppression in only 1% (n = 2 of 187 neurons)(Fig. 2d,e). The paucity of suppressive responses was unlikely to result from low spontaneous activity under anesthesia, as a similarly low proportion of suppressive responses was observed in awake mice, despite their significantly higher spontaneous firing rates (Supplementary Fig. 4).

Excitatory responses were robust across trials and remained stable across repeated pulses in a 5-Hz stimulus train (Fig. 2f). Prolonged stimulation (200 ms) elicited sustained responses that gradually adapted over the stimulation period (Fig. 2g,h), indicating that the corticocollicular pathways can reliably transmit sustained cortical input to the IC.

To assess whether these responses were consistent with monosynaptic connectivity, we analyzed first-spike latencies after stimulus onset. Because response latencies decreased with increasing laser intensity (Supplementary Fig. 3e-h), relatively high laser intensities were used for this analysis. First-spike latencies varied widely across IC neurons, ranging from 4.4 to 52.8 ms for S1 stimulation (median: 19.4 ms, n = 113 neurons) and from 5.97 to 56.8 ms for M1 stimulation (median: 22.3 ms, n = 68 neurons), with no significant difference between the two pathways (Fig. 2i,j; Wilcoxon rank-sum test, p = 0.09). The broad latency distributions suggest that S1- and M1-evoked responses reflect a mixture of monosynaptic and polysynaptic inputs, consistent with previous findings using electrical stimulation in rats^34^.

We next examined the convergence of S1 and M1 inputs onto individual IC neurons by recording responses to stimulation of both pathways (n = 68 responsive neurons). The majority of these neurons (59%) responded to both S1 and M1 stimulation, whereas 29% responded selectively to S1 and 12% selectively to M1 (Fig. 2k). Among dual-responsive neurons, S1 stimulation evoked significantly stronger responses than M1 stimulation (Fig. 2l; median firing rate: S1, 24.1 Hz; M1, 12.5 Hz; n = 40, p = 0.021). Responses in S1- and M1-selective neurons were weaker than those in dual responsive neurons (S1 dual vs S1 selective, p = 0.009; M1 dual vs M1 selective, p = 0.0019) and did not differ significantly from each other (S1 selective vs M1 selective, p = 0.27; Wilcoxon rank-sum tests).

Reconstruction of recording sites revealed that S1- and M1-responsive neurons were more abundant in rostral IC sections, with relatively few responsive neurons detected at the most caudal level (Fig 2m,n). Responsive neurons were distributed across all major IC subdivisions, including the CNIC, but were most prevalent in the LCIC (Fig. 2o). This spatial distribution was broadly consistent with the transsynaptic tracing results (Fig. 1e-h), which similarly showed that cortical recipient neurons were most abundant in the rostral IC and concentrated in the LCIC.

### S1 and M1 inputs recruit both auditory and non-auditory IC neurons

To understand how auditory inputs interact with S1 and M1 inputs, we first examined how individual IC neurons responded to sound (50 ms broadband noise, 70 dB SPL) and cortical stimulation. We identified neurons that responded to both stimuli, to cortical stimulation alone, or to sound alone (Fig. 3a). Given the IC’s central role in auditory processing, we expected most cortical-responsive neurons to also respond to sound. Surprisingly, among S1-responsive IC neurons, 58% did not respond to noise, and among M1-responsvie IC neurons, 63% did not respond (Fig. 3b,c). To determine whether these neurons might respond to other sounds, we additionally presented pure tones across a range of frequencies. Even when probed with both stimulus types, 50% of S1-responsive and 57% of M1-responsive neurons remained unresponsive to auditory stimulation (Fig. 3b,c). Thus, a substantial population of IC neurons receiving functional input from S1 or M1 is not driven by broadband noise or pure tones.

**Figure 3.**
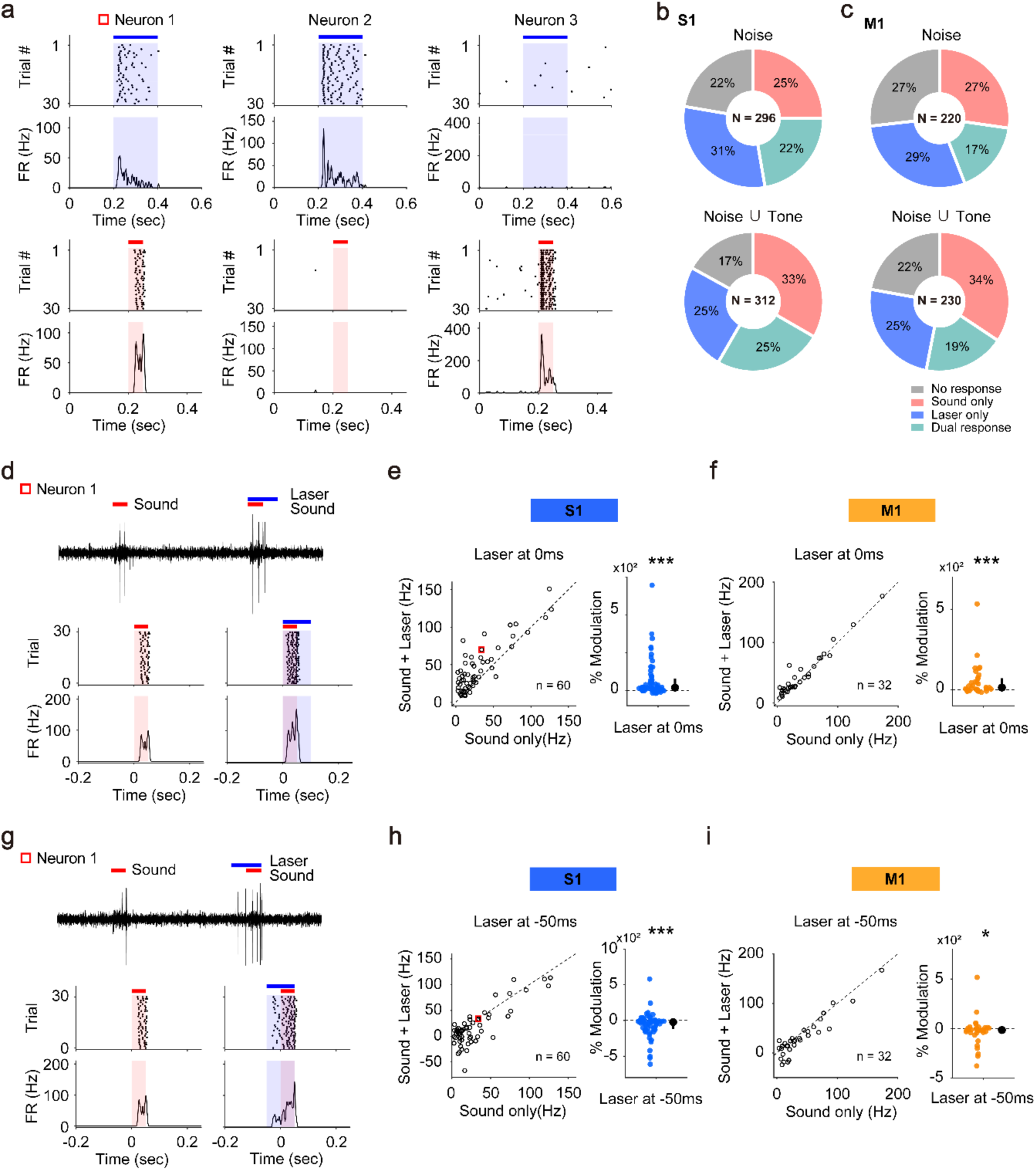
Modulation of sound-evoked responses in IC neurons by S1 and M1 corticocollicular inputs. **a**) Example IC neurons responsive to both sound and S1 stimulation (Neuron 1), S1 stimulation only (Neuron 2), or sound only (Neuron 3). Raster plots and PSTHs are shown. Blue bars and shading indicate laser stimulation (200 ms); red bars and shading indicate sound presentation (50 ms). **b-c**) Proportions of IC neurons responsive only to sound, only to cortical stimulation, to both, or to neither during S1 (**b**) or M1 (**c**) stimulation. Responses are shown separately for broadband noise (top), and responses to either broadband noise or pure tones combined (bottom). **d-f**) Effects of S1 (**e**) and M1 (**f**) stimulation on sound-evoked responses when laser stimulation began at sound onset. **d**) Example IC neuron showing responses to sound alone and to concurrent sound and S1 stimulation. Raw spike waveforms (top), raster plots (middle), and PSTHs (bottom) are shown. Red bars and shading indicate sound presentation; blue bars and shading indicate laser stimulation. **e-f**) Scatter plots (left) compare sound-evoked firing rates with and without cortical stimulation, and plots on the right show percent modulation of sound-evoked responses (**e**, n = 60, p = 1.21 × 10^-8^; **f**, n = 32, p = 5.80 × 10^-4^, Wilcoxon signed-rank test). The red square indicates the example neuron shown in (**d**). Blue and orange circles represent individual neurons for S1 (**e**) and M1 (**f**), respectively, and black circles and error bars indicate the median and interquartile range of the population**. g-i**) Corresponding analyses for laser stimulation beginning 50 ms before sound onset (**h**, n = 60, p = 2.84 × 10^-4^; **i**, n = 32, p = 0.0167; Wilcoxon signed-rank tests).

We next investigated how S1 and M1 inputs influence auditory responses in sound-responsive IC neurons that also exhibited excitatory responses to cortical stimulation (S1, 60 of 140 neurons; M1, 32 of 97 neurons). When cortical stimulation began concurrently with sound, sound-evoked responses were significantly enhanced relative to sound alone (S1: median modulation, 40%, Wilcoxon signed-rank test, p = 1.21 × 10^-8^; M1: median modulation, 14%, p = 5.80 × 10^-4^; Fig. 3d-f). In contrast, when cortical stimulation preceded sound onset by 50 ms, sound-evoked responses, measured relative to the baseline activity immediately preceding sound onset^47^, were significantly suppressed (S1: median modulation, −23%, p = 2.84 × 10^-4^; M1: median modulation, −16%, p = 0.0167; Fig. 3g-i). Thus, the influence of S1 and M1 inputs on auditory responses depended on their timing relative to sound, with coincident cortical activity enhancing sound-evoked responses and preceding cortical activity suppressing them.

### S1 and M1 inputs modulate frequency tuning in the IC

To further examine whether S1 and M1 inputs influence sound frequency processing, we recorded responses to pure tones (2-64 kHz, 50 ms, 70dB SPL) in S1- and M1-responsive IC neurons while optogenetically activating IC-projecting cortical neurons. We used two stimulation protocols to probe the effects of non-auditory cortical input: “transient” stimulation, in which a laser pulse began 50 ms before tone onset (Fig. 4a), and “sustained” stimulation, in which a ramped laser pulse^57^ began 300 ms before tone onset to mimic prolonged cortical drive (Fig. 4b).

**Figure 4.**
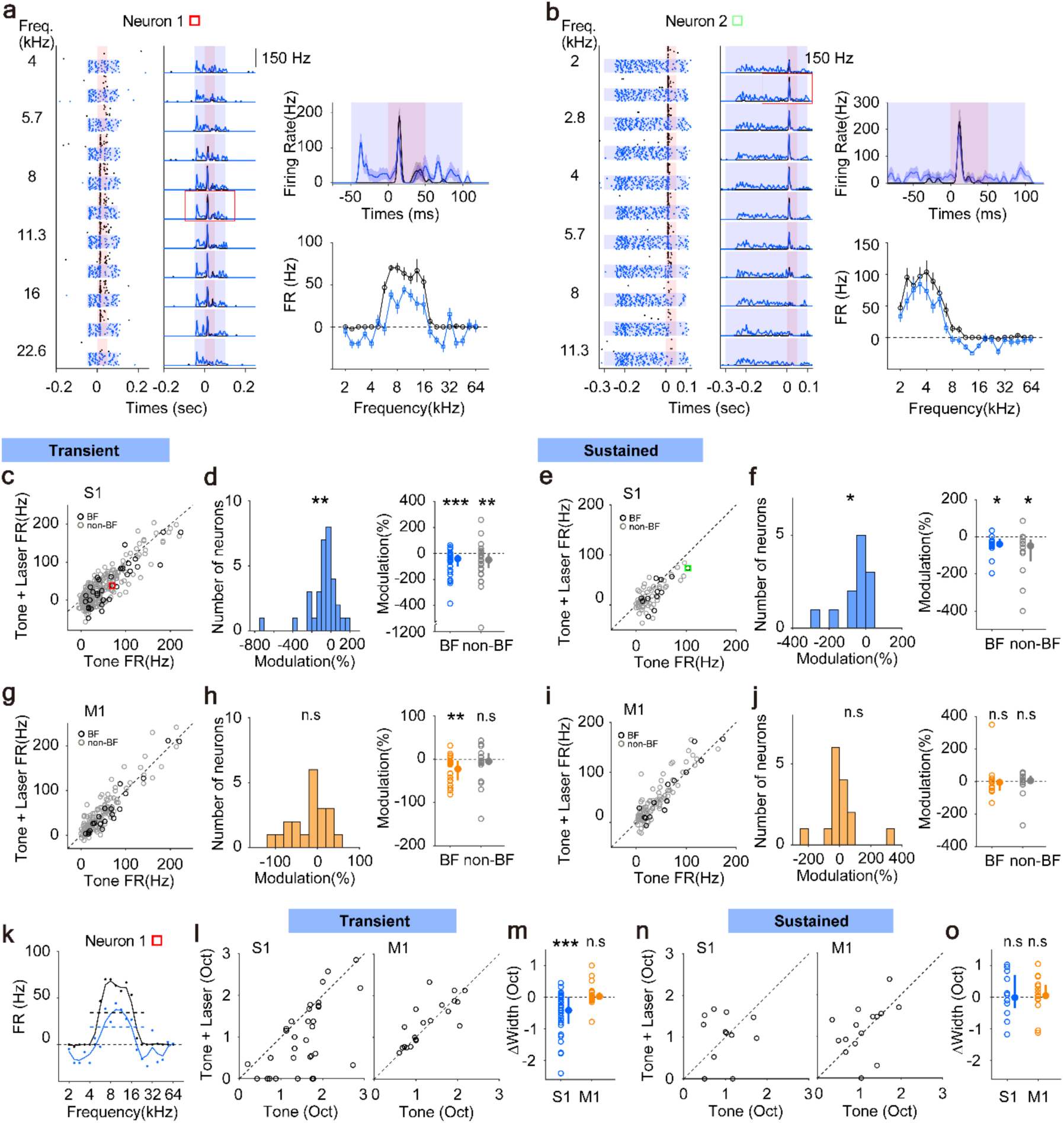
Modulation of tone-evoked responses and frequency tuning in the IC by S1 and M1 corticocollicular inputs. **a-b**) Responses of two example IC neurons to pure tones with (blue) or without (black) optogenetic S1 stimulation. Transient (**a**) and sustained (**b**) stimulation protocols are shown. *Left:* Raster plots and PSTHs across tone frequencies. The red box indicates the best frequency (BF). *Right:* PSTHs at the BF (top) and frequency tuning curves (bottom). Red and blue shading indicate the periods of tone and laser stimulation, respectively. **c-f**) Effects of S1 stimulation on tone-evoked responses. **c,e**) Scatter plots comparing tone-evoked firing rates with and without S1 stimulation. Gray circles represent non-BF responses, and black circles represent BF responses. The red and green squares indicate the example neurons shown in (**a**) and (**b**), respectively. Dashed lines indicate unity. **d,f**) Distribution of response modulation (left) and modulation of BF and non-BF responses (right). Transient S1 stimulation significantly shifted the distribution of response modulation (p = 0.0031) and suppressed both BF (p = 3.81 x 10^-4^) and non-BF responses (p = 0.006). Sustained S1 stimulation similarly shifted the distribution of response modulation (p = 0.01) and suppressed both BF (p = 0.021) and non-BF responses (p = 0.019; Wilcoxon signed-rank tests). **g-j**) Same as (**c-f**), but for M1 stimulation. Transient M1 stimulation did not significantly shift the overall distribution of response modulation (p = 0.093) but significantly suppressed BF responses (p = 0.0032), with no significant effect at non-BF frequencies (p = 0.396). Sustained M1 stimulation did not significantly affect the overall distribution of response modulation (p = 0.89), BF responses (p = 0.24), or non-BF responses (p = 0.68; Wilcoxon signed-rank tests). **k**) Smoothed frequency tuning curves for the example neuron shown in (**a**), with (blue) and without (black) S1 stimulation. Tuning width was quantified as the full width at half maximum (horizontal dashed line). **l**) Scatter plots comparing tuning width measured with and without S1 or M1 stimulation during transient stimulation. **m**) Population summary of changes in tuning width during transient cortical stimulation (S1, p = 1.43 x 10^-4^; M1, p = 0.48). **n**) Scatter plots comparing tuning width measured with and without S1 or M1 stimulation during sustained stimulation. **o**) Population summary of changes in tuning width during sustained cortical stimulation (S1, p = 0.89; M1, p = 0.41; Wilcoxon signed-rank tests).

Tone-evoked responses, measured relative to the baseline activity immediately preceding tone onset, were significantly attenuated during S1 stimulation compared with tone presentation alone (Fig. 4c-f; transient: median modulation, -48%, n = 32, p = 0.0031; sustained: median modulation, -38%, n = 12, p = 0.0137; Wilcoxon signed-rank tests). Attenuation was evident at both best frequencies (BFs), where tone-evoked responses were largest (transient: BF, p = 3.81 x 10^-4^; non-BF, p = 0.006; sustained: BF, p = 0.021; non-BF, p = 0.019). In contrast, attenuation during M1 stimulation was weaker and reached significance only at BFs during transient stimulation (Fig. 4g-j; transient: median modulation, -5.0%, n = 20, p = 0.093; BF: p = 0.0032; non-BF: p = 0.396; sustained: median modulation, -4.6%, n = 15, p = 0.89; BF: p = 0.24; non-BF: p = 0.68; Wilcoxon signed-rank tests).

We next asked whether cortical stimulation alters frequency selectivity. Tuning width was quantified as the full width at half maximum of smoothed tuning curves (Fig. 4k). Transient S1 stimulation significantly decreased tuning widths, indicating increased frequency selectivity (median Δwidth, -0.4 octaves, n = 32; p = 1.43 x 10^-4^, Wilcoxon signed-rank test), whereas transient M1 stimulation did not significantly alter tuning width (median Δwidth, 0.25 octaves, n = 20, p = 0.48; Fig. 4l,m). Sustained stimulation produced heterogeneous effects across neurons, with no significant population level changes in tuning width for either S1 (median Δwidth, -0.013 octaves, n = 12, p = 0.89) or M1 stimulation (median Δwidth, 0.05 octaves, n = 15, p = 0.41; Fig. 4n,o).

Together, these findings show that S1 and M1 corticocollicular inputs differentially modulate frequency processing in the IC. Transient S1 stimulation produced the most prominent effects, attenuating tone-evoked responses and narrowing frequency tuning, whereas M1 and sustained stimulation produce weaker and more variable changes.

### IC responses to cortical stimulation correlate with locomotion-induced modulation

Recent evidence indicates that neural activity in the IC is strongly influenced by movement, including locomotion. While these findings suggest that the IC integrates auditory and movement-related information, the sources of these locomotion-related signals remain poorly understood. Because both the somatosensory and motor cortices are strongly engaged during locomotion^58^, we asked whether S1 and M1 projections to the IC contribute to locomotion-related modulation.

To address this question, we optogenetically activated IC-projecting S1 or M1 neurons in awake, head-fixed mice during spontaneous locomotion (Fig. 5a). We first identified IC neurons responsive to cortical stimulation (Fig. 5b) and then examined whether these same neurons were also modulated during locomotion (Fig. 5c). The example neuron shown was excited by both S1 stimulation and locomotion (Fig. 5b,c). Across the population, 88% of cortical-responsive neurons (38 of 43) were significantly modulated during locomotion (Fig. 5c,d), with 79% (30 of 38) showing excitation and 21% (8 of 38) showing suppression (Fig. 5e-g). Moreover, the magnitude of locomotion-induced modulation was positively correlated with responsiveness to cortical stimulation. Specifically, neurons exhibiting stronger responses to S1 or M1 activation also showed larger locomotion-related modulation (Spearman’s ρ = 0.47, p = 0.012; Fig. 5h). These findings indicate a close relationship between descending sensorimotor cortical input and locomotion-related activity in IC neurons.

**Figure 5.**
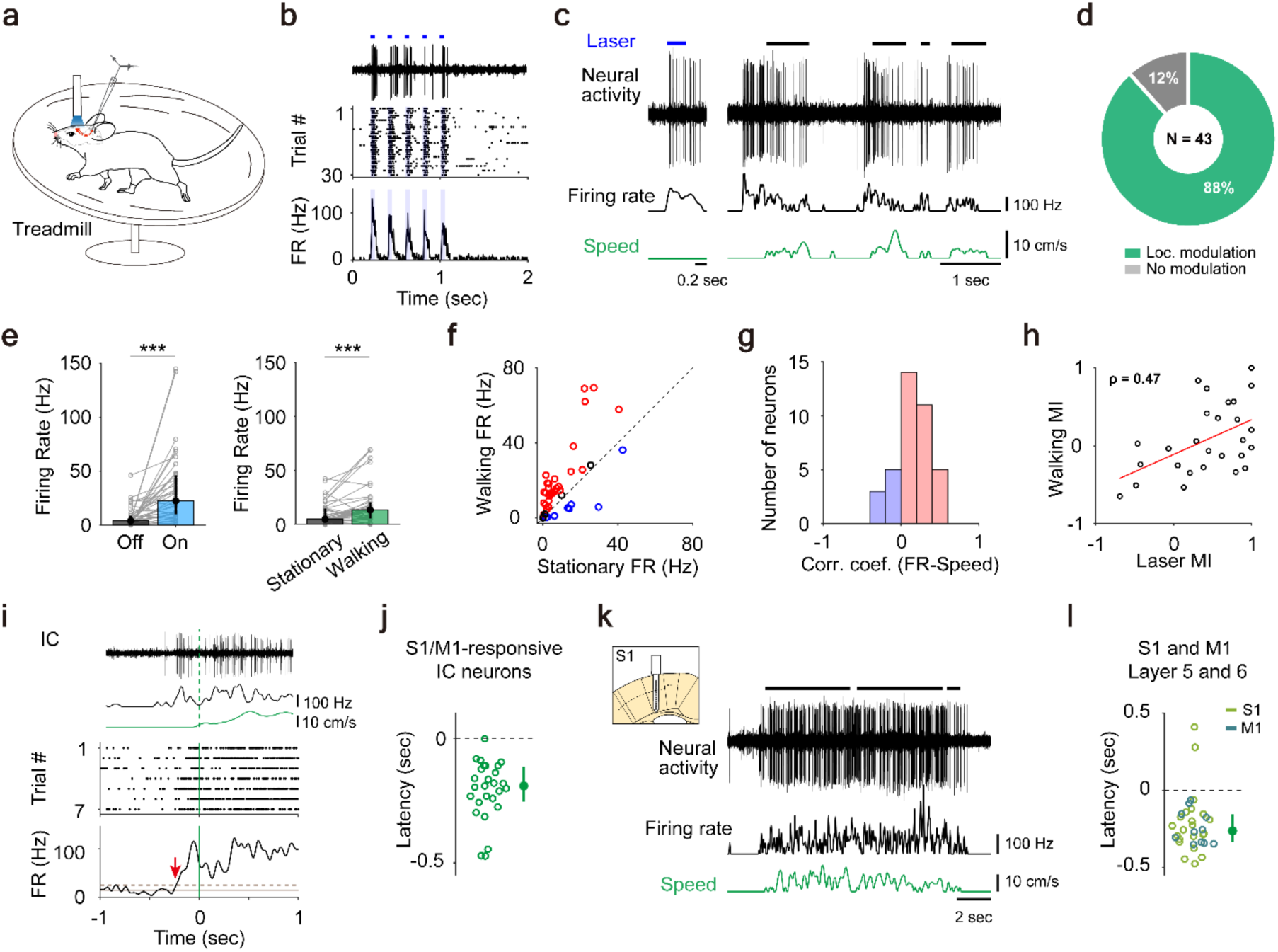
Relationship between S1 and M1 corticocollicular inputs and locomotion-related modulation in IC neurons. **a**) Schematic of the experimental setup. IC recordings and optogenetic stimulation of corticocollicular neurons were performed in awake, head-fixed mice running on a circular treadmill. **b-c**) Example IC neuron showing responses to S1 stimulation (**b**) and modulation during locomotion (**c**). **b**) Spike waveform, raster plot, and PSTH during optogenetic stimulation. Blue horizontal bars indicate laser stimulation. **c**) Neural activity, smoothed firing rate, locomotion speed (green). Black horizontal bars indicate locomotion periods. **d**) Proportion of locomotion-modulated neurons among S1- and M1-responsive IC neurons (n = 43). **e**) Population data showing firing rates during laser stimulation (left) and locomotion (right) (Wilcoxon signed-rank test; laser, p = 1.53 x 10^-6^; locomotion, p = 3.14 x 10^-4^). **f**) Comparison of firing rates during stationary and walking periods in S1- and M1-responsive IC neurons. Neurons are color-coded according to the direction of locomotion-related modulation: increased (red), decreased (blue), or not significantly modulated (black). **g**) Distribution of correlation coefficients between firing rate and locomotion speed. Red: increased neurons; blue: decreased neurons. **h**) Relationship between modulation indices (MI) for laser stimulation and locomotion (Spearman’s ρ = 0.47, p = 0.012). **i**) Example S1-responsive IC neuron exhibiting increased firing before locomotion onset. Top: neural activity, smoothed firing rate, and locomotion speed. Botton: raster plot and PSTH aligned to locomotion onset. Green vertical line: locomotion onset; red arrow: the onset of neural modulation. Solid horizontal line indicates the baseline firing rate; dashed horizontal line indicates the threshold (2 SD above baseline) used to determine modulation onset. **j**) Distribution of modulation onset latencies relative to locomotion onset for S1- and M1-responsive IC neurons (n = 27). **k**) Example recording from a layer 5 S1 neuron showing locomotion-related modulation. **l**) Modulation onset latencies relative to locomotion onset for deep-layer S1 and M1 neurons (S1, n = 23; M1, n = 11).

Previous studies have reported that some locomotion-modulated IC neurons change their activity before movement onset^29,30^. We therefore examined the temporal relationship between neural activity and locomotion onset in cortical-responsive IC neurons. Many of these neurons exhibited significant changes in firing rate before movement onset, as illustrated by the example neuron in Fig. 5i. Across the population, the onset of neural modulation preceded locomotion onset, with a median latency of -192 ms (Fig. 5j; n = 27 neurons).

To determine whether corticocollicular neurons exhibit similar anticipatory activity, we next recorded from putative layer 5 and 6 neurons in S1 and M1. As a population, these neurons also exhibited anticipatory modulation before locomotion onset (Fig. 5k,l; median latency = -262.5 ms, n = 34 neurons). Consistent with this result, light-responsive cortical multiunit clusters, which included IC-projecting neurons, also exhibited anticipatory activity (Supplementary Fig. 5; median latency = -207 ms).

Together, these findings show that IC neurons receiving functional input from S1 and M1 are strongly modulated during locomotion. The observation that anticipatory activity in both cortical-responsive IC neurons and sensorimotor cortical neurons exhibit locomotion-related modulation and anticipatory activity suggests that descending corticocollicular pathways provide a source of movement-related signals to the IC.

## Discussion

In this study, we investigated how somatosensory and motor corticocollicular projections contribute to sensorimotor integration in the auditory midbrain. We found that trunk- and limb-related regions of S1 and M1 provide prominent monosynaptic input to the IC, and that optogenetic activation of these projections robustly drove IC neural activity. Approximately half of the cortical-responsive IC neurons were unresponsive to broadband noise or pure tones, revealing a distinct population of IC neurons preferentially driven by descending sensorimotor input. In sound-responsive neurons, S1 and M1 activation bidirectionally modulated auditory responses depending on the timing of cortical activation relative to sound. Finally, both cortical-responsive IC neurons and deep-layer S1 and M1 neurons exhibited locomotion-related modulation and anticipatory activity before movement onset, suggesting that descending corticocollicular pathways convey movement-related signals to the IC. Together, these findings identify a descending sensorimotor circuit that integrates body- and movement-related information with auditory processing in the midbrain.

### Functional organization of S1 and M1 projections to the IC

Our anatomical tracing revealed that corticocollicular projections from S1 and M1 arise predominantly from trunk- and limb-related subregions. Although projections from S1 and M1 to the IC have been described previously^31,32,34,45^, their relative somatotopic contributions have remained unclear. Prior work has focused primarily on whisker-related S1 projections, which preferentially target the IC shell and modulate auditory processing^21,33^. In contrast, our tracing revealed denser projections from trunk and limb regions than from whisker S1. Our findings raise the question of how whisker- and body-related cortical regions contribute complementary information about the environment and body state during exploratory behavior.

Transsynaptic tracing further demonstrated that both S1 and M1 establish direct monosynaptic connections with IC neurons. Approximately 70% of recipient neurons were located within the IC shell, consistent with the concentration of corticocollicular terminals in the LCIC^31^, whereas nearly one-third were located in the CNIC, indicating that direct non-auditory cortical influences extend into the lemniscal auditory pathway^34^. Most recipient neurons were GAD-negative, similar to targets of auditory corticocollicular projections^56,59,60^. In contrast, whisker-related S1 inputs preferentially target GABAergic neurons in the IC shell^33,61^. Together, these findings suggest that distinct somatotopic corticocollicular pathways engage different IC circuit elements and may therefore influence auditory processing through distinct mechanisms.

Optogenetic activation of IC-projecting S1 and M1 neurons predominantly evoked excitation, whereas suppression was observed in only a small fraction of responsive neurons.

This predominance of excitation is consistent with anatomical evidence that corticocollicular neurons arising from layers 5 and 6 are glutamatergic^59,62,63^, as well as physiological studies demonstrating excitatory corticocollicular inputs^34,47,54,55^. However, the broad distribution of response latencies indicates that cortical signals are further processed by local polysynaptic circuits^64,65^. The scarcity of suppressive responses further suggests that local GABAergic circuits are recruited less often by S1 and M1 projections, making feedforward inhibition relatively uncommon^56^. This interpretation is further supported by our anatomical finding that most cortical recipient neurons are non-GABAergic.

Previous work suggested that diverse cortical inputs converge onto overlapping IC neuron populations^34^. Consistent with this idea, approximately 60% of responsive IC neurons in our recordings responded to both S1 and M1 stimulation. Because somatosensory and motor cortical activity are tightly coordinated during movement, convergent corticocollicular inputs may allow IC neurons to integrate information related to body position, movement, and sensory consequences of actions^61^. Similar convergence has been described in the superior colliculus^66^ and higher order thalamus^67^, suggesting that integration of descending somatosensory and motor cortical signals may represent a common organizational principle of subcortical sensorimotor circuits.

### Modulation of auditory processing by non-auditory cortical inputs

Our results demonstrate that the influence of non-auditory corticocollicular inputs on auditory processing depends critically on stimulus timing. When S1 and M1 activation coincided with sound presentation, auditory responses were enhanced across IC subregions, potentially facilitating detection of behaviorally relevant events accompanied by concurrent movement or somatosensory signals. In contrast, when cortical activation preceded sound presentation and established an elevated baseline level of activity, auditory responses were attenuated and frequency tuning became narrower. These effects were strongest during S1 stimulation and during transient, compared with sustained, cortical activation.

Previous studies have reported both enhancement and suppression of auditory responses following somatosensory stimulation, depending on the pathway activated. Whisker stimulation produces both enhancement and suppression in the LCIC, but little modulation in the CNIC^21,33^. Activation of ascending somatosensory pathways often suppresses auditory activity^38,40^, whereas high frequency vibration enhances auditory responses in the LCIC^22^. Together with our findings, these studies suggest that the influence of somatosensory inputs depend both on the anatomical pathways engaged and on the temporal relationship between somatosensory and auditory signals.

Enhancement and attenuation are likely to serve complementary rather than opposing functions. For example, coincident corticocollicular inputs may increase the salience of multisensory events during movement, whereas preceding cortical activity may suppress predictable and self-generated auditory inputs while preserving auditory sensitivity along with higher frequency selectivity^29,68,69^. Together, these mechanisms could enhance the detection of unexpected or behaviorally relevant sounds during active behaviors. Beyond sensory gating, non-auditory modulation may also support broader functions such as dynamic allocation of processing resources^12,61^ or simultaneous processing of sensory and behavioral information^16^. Thus, the functional consequences of corticocollicular modulation are likely to depend on behavioral context. Regardless of the specific role, our findings demonstrate that descending somatosensory and motor cortical signals dynamically regulate auditory processing at the level of the midbrain.

### Potential roles of S1 and M1 inputs during movement

Approximately half of S1- and M1-responsive IC neurons were unresponsive to broadband noise or pure tones. Although some of these neurons may respond selectively to more complex acoustic stimuli or exhibit only subthreshold auditory responses, previous studies have also reported IC neurons that respond preferentially to somatosensory stimulation^21,22,38^. Our results therefore suggest that a substantial population of IC neurons may primarily encode behavioral context rather than acoustic stimuli. This interpretation is consistent with growing evidence that IC neurons encode behavioral state, task, and movement variables in addition to sensory inputs^45,70–74^. Together with descending inputs from the auditory cortex^75^, S1 and M1 projections may contribute to a broader representation of behavioral information within the IC.

Prior work has shown that locomotion robustly modulates IC activity and that many IC neurons exhibit changes in firing before movement onset^29^. Moreover, this modulation persists in deaf mice, indicating that movement-related signals are generated centrally^30^. Although ascending somatosensory pathways likely contribute to locomotion-related modulation^22,76^, they cannot readily explain the pre-movement activity observed in many IC neurons.

Several observations suggest that descending corticocollicular pathways provide a source of movement-related signals in the IC. First, the large majority of S1- and M1-responsive IC neurons were significantly modulated during locomotion, with many neurons exhibiting anticipatory activity before movement onset. Second, the magnitude of locomotion-related modulation correlated with the strength of S1- and M1-evoked responses, suggesting that cortical responsiveness and locomotion-related activity arise, at least in part, from shared descending inputs. Finally, deep-layer S1 and M1 neurons, which include corticocollicular neurons, were themselves strongly modulated during locomotion and exhibited anticipatory activity. The presence of pre-movement activity in S1 is consistent with recent evidence that somatosensory cortex participates in motor control and exhibits pre-movement activity alongside motor cortex rather than simply encoding somatosensory feedback^58,77^.

Although these observations support a role for corticocollicular pathways in conveying movement-related signals, several features of locomotion-related modulation were not reproduced by S1 and M1 stimulation. Suppression of IC neural activity was commonly observed during locomotion but was rarely evoked by optogenetic activation of S1 and M1 inputs. Although excitatory cortical inputs could produce suppression through recruitment of local IC circuits^56^, the scarcity of suppressive responses suggests that additional inputs contribute to locomotion-related suppression. Likewise, locomotion has been reported to sharpen frequency tuning in IC neurons^29^. Although transient S1 stimulation similarly sharpened frequency tuning, this effect was not observed during sustained S1 stimulation or during M1 stimulation. Together, these findings indicate that descending S1 and M1 pathways constitute one component of a broader network of inputs that shapes locomotion-related modulation in the IC.

Recent studies increasingly indicate that the IC integrates behavioral, motor, and sensory information. Our findings identify a descending sensorimotor circuit that contributes to this integration. During natural behavior, such a circuit may allow auditory processing to be continuously adjusted according to ongoing movement and behavioral context, enabling the auditory midbrain to represent sounds in relation to the animal’s own actions.

## Methods

### Animals

All experimental procedures were approved by the Institutional Animal Care and Use Committee of the Korea Brain Research Institute (IACUC-23-00077-M2). Mice were group-housed under a 12 h light/12 h dark cycle with ad libitum access to food and water. Wild-type and transgenic mice of both sexes were used and were 6-8 weeks old at the time of surgery. C57BL/6J mice (JAX #000664) were used for optogenetic experiments, including functional characterization of corticocollicular projections, assessment of their effects on auditory responses under anesthesia, and examination of locomotion-related modulation in awake mice. Ai9 (JAX #007909) and Ai96 (JAX #028866) transgenic mice were used for anterograde and retrograde anatomical tracing, respectively.

### Viral vectors

For retrograde tracing of cortical inputs to the IC, AAV-retro-hSyn-Cre-P2A-tdTomato (Addgene #107738-AAVrg) was injected into the IC of Ai96 mice. For anterograde transsynaptic tracing from the cortex, a self-complementary version of AAV1-hSyn-Cre-WPRE-hGH (VectorBuilder) was injected into the cortex of Ai9 mice. For optogenetic manipulation of corticocollicular neurons, retrograde AAVs encoding channelrhodopsin-2 (AAV-retro-hSyn-hChR2(H134R)-EYFP (Addgene #26973-AAVrg) or AAV-retro-CAG-hChR2(H134R)-tdTomato (Addgene #28017-AAVrg)) were injected into the IC.

### Stereotaxic viral injection

Mice were placed in a stereotaxic frame and anesthetized with isoflurane delivered via a nose cone, with room air as the carrier gas (2-4% for induction and 1.5-2% for maintenance). The depth of anesthesia was periodically assessed by the toe-pinch reflex. Eye ointment was applied to prevent corneal drying. Lidocaine was administered subcutaneously under the scalp for local anesthesia, followed by an incision to expose the skull. For IC injections, a small craniotomy, ∼1.5 mm in diameter, was made over the IC (AP, -5.0 mm relative to bregma; ML, 1.2 mm from the midline). For cortical injections, a craniotomy smaller than 1 mm in diameter was made over S1 (AP, -0.3 mm; ML, 2.3 mm) or M1 (AP, -0.3 mm; ML, 1.0 mm). Viral vectors were loaded into a glass pipette (20 µm inner tip diameter) and injected at two depths, 1.0 and 0.5 mm below the pial surface for both IC and cortical injections, using a syringe pump at a rate of 80 nL/min. The injection volume was 150 nL at each depth for the IC and 100 nL at each depth for the cortex. After each injection, the pipette was left in place for 15 min to allow diffusion and then slowly withdrawn. The incision was sutured, and ketoprofen (5 mg/kg) was administered subcutaneously for postoperative analgesia. Mice were allowed to recover on a heating pad before being returned to their home cages. Experiments were performed 8-10 weeks later to ensure robust viral expression.

### Headpost implantation

Mice were anesthetized with isoflurane using the same procedures as described above for stereotaxic viral injection. An incision was made to expose the skull, and the skull surface was cleaned and dried. A custom-made headpost was affixed to the skull using dental cement (C&B Metabond). Ketoprofen (5 mg/kg) was administered subcutaneously for postoperative analgesia, and mice were allowed to recover on a heating pad before being returned to their home cages. Animals were monitored daily for signs of infection or other postoperative complications. For electrophysiological recordings in awake mice, animals were habituated to head fixation and walking on a circular treadmill for at least 1 week through daily 30-min sessions.

### Electrophysiological recording

For anesthetized recordings, mice were anesthetized with an intraperitoneal injection of ketamine/xylazine (100 and 10 mg/kg, respectively), with supplemental doses (25 and 1.25 mg/kg, respectively) administered as needed to maintain anesthesia. Body temperature was maintained at 37°C, and the depth of anesthesia was monitored by the toe-pinch reflex and the absence of spontaneous whisking. Eye ointment was applied to prevent corneal drying. For IC recordings, a ∼2 mm diameter cranial window was made over the IC. For optogenetic stimulation of cortical neurons, a cranial window approximately 2.5 × 2 mm in size were made over the S1 and M1 cortical regions. Extracellular recordings from the IC were performed using a linear array of tungsten electrodes (∼5 MΩ, 200 μm spacing; FHC, Bowdoin, ME, USA) mounted on a motorized micromanipulator (IVM Mini, Scientifica, Uckfield, UK). Neural signals were acquired using a 16-channel headstage (RHD2132; Intan Technologies, Los Angeles, CA, USA) and an Open Ephys data acquisition system (Open Ephys GUI v.0.5.5). Spiking activity was band-pass filtered at 600-6000 Hz and digitized at 30 kHz.

For awake recordings, cranial windows over the IC and target cortical regions were made under isoflurane anesthesia and sealed with Kwik-Cast (WPI, Sarasota, FL, USA). Ketoprofen (5 mg/kg) was administered subcutaneously for analgesia. Mice were allowed to recover for at least 3 h before recording, with most recordings performed the following day. Extracellular recordings were performed using the same procedures as for anesthetized recordings. Locomotion on the treadmill was monitored using a rotary encoder (Scitech Korea, Korea), and the encoder output was recorded with neural signals for offline analysis^78^.

### Acoustic stimuli

Acoustic stimuli were generated in MATLAB (MathWorks, Natick, MA, USA) at a sampling rate of 250 kHz. Sound stimuli were delivered via a digital-to-analog converter (PCIe-6343, National Instruments), amplified with a power amplifier (#70103, Avisoft, Glienicke/Nordbahn, Germany), and presented through an ultrasonic speaker (Vifa, Avisoft). The speaker was positioned 15 cm from the animal’s right ear at a 45° angle relative to the midline. The sound system was calibrated using a ¼-inch microphone (Brüel & Kjær 4939, Nærum, Denmark). Stimulus presentation was controlled by a custom-written Python program (Kranky; https://bitbucket.org/spikeCoder/kranky) interfaced with the Open Ephys GUI software. Broadband noise spanning 2-64kHz was presented at 70 dB SPL for 50 ms, with 5ms onset and offset ramps. Pure tones ranging from 2 to 64kHz in quarter-octave steps were presented at 70 dB SPL for 50 ms, with 1 ms onset and offset ramps, in pseudorandom order. Each frequency was presented at least 20 times.

### Optogenetic stimulation

IC-projecting cortical neurons were optogenetically stimulated using a blue laser (473 nm). Light was delivered through an optical fiber with a 200 µm core diameter (Thorlabs) positioned on the dural surface above the target cortical area. For most experiments, laser power was set to 40-50 mW at the fiber tip. To examine the effects of cortical activation on IC activity, laser stimulation was delivered either as a single 200 ms pulse or as a train of five 50 ms pulses at 5 Hz. Each trial lasted 2 s, and each stimulus condition was repeated at least 30 times. To assess modulation of noise-evoked responses, a 100 ms laser pulse was delivered either simultaneously with the noise stimulus or beginning 50 ms before noise onset. To assess modulation of tone-evoked responses, two stimulation protocols were used: a transient protocol consisting of 150 ms square laser pulse beginning 50 ms before tone onset, and a sustained protocol consisting of 400 ms laser pulse with a 100 ms linear onset ramp to provide sustained cortical activation. In the sustained protocol, the tone was presented 300 ms after laser onset.

### Histology

At the end of each experiment, recording sites were marked by applying current pulses (30 µA, 10 s) through the recording electrodes at three depths: 500, 1000, and 1500 µm below the pial surface. Animals were then transcardially perfused with saline followed by 4% paraformaldehyde. Brains were post-fixed overnight, cryoprotected in 30% sucrose, embedded in OCT compound, and sectioned coronally at 40 µm on a cryostat (CM1860, Leica). For GAD67 immunostaining, free-floating sections were rinsed in 0.1 M phosphate-buffered saline (PBS) and blocked for 2 h in PBS containing 0.1% Triton X-100, 2% normal goat serum, and 1% bovine serum albumin. Sections were then incubated overnight at 4°C with a mouse anti-GAD67 antibody (1:1000; Sigma, Cat. #MAB5406), rinsed in PBS, and incubated with an Alexa Fluor 488-conjugated goat anti-mouse IgG secondary antibody (1:2000; ThermoFisher, Cat. #A11001). Images were acquired using a Leica TCS SP8 confocal microscope (Brain Research Core Facilities at KBRI) with a 40 x water-immersion objective.

### 3D anatomical reconstruction

Thirty coronal brain sections, each 40 µm thick and sampled at 120 µm intervals, were imaged using an inverted microscope with a 4 x objective. Images were processed with AMaSiNe software^52^ for automated alignment to the Allen Mouse Brain Reference Atlas and 3D reconstruction and visualization of IC-projecting cortical neurons.

### Cell counting

For quantification of cortical-recipient and GAD67-positive IC neurons, four mice were analyzed: two with S1 injections and two with M1 injections. Confocal images were acquired from three IC sections per mouse, spanning the rostrocaudal axis, and processed using ImageJ. Fluorescently labeled neurons were manually counted and cortical-recipient neurons co-labeled for GAD67 were classified as GAD67+, whereas those lacking GAD67 immunoreactivity were classified as GAD67-. The proportions of GAD67+ and GAD67-cortical-recipient neurons were calculated across the rostrocaudal axis and IC subdivisions for each mouse. Data from S1 and M1 injections were pooled for analysis.

### Neural data analysis

For single unit spike sorting, spikes were detected and sorted offline using Offline Sorter v4 (Plexon, Dallas, TX, USA). Spike waveforms were clustered using principal component analysis. Well-isolated clusters that were significantly separated from other clusters (p < 0.05, multivariate analysis of variance) and had fewer than 0.5% of spikes violating a 0.7 ms refractory period were classified as single units and included in subsequent analyses.

Laser-evoked activity was quantified as the mean firing rate within a response window beginning at stimulus onset (0∼210 ms for single pulse or 0∼60 ms for each pulse during pulse train stimulation), relative to spontaneous activity measured during a 100 ms baseline window (-130 to -30 ms relative to laser onset). For each neuron, the significance of laser-evoked responses was determined by comparing firing rates during the baseline and response periods across trials (p < 0.05, paired t-test). PSTHs were constructed for each stimulus condition and smoothed with a Gaussian kernel (σ = 2 ms). First-spike latency was defined as the time from laser onset to the first subsequent evoked spike.

Broadband noise-evoked responses were quantified as the mean firing rate during the 50 ms sound presentation relative to the baseline firing rate measured during 100 ms window preceding sound onset. For trials in which laser stimulation preceded sound onset, the 20-ms laser-only period immediately preceding sound onset served as the baseline. For each neuron, the significance of the sound-evoked response was determined by comparing firing rates during the baseline and sound periods across trials (p < 0.05, paired t-test). Similarly, the significance of laser-evoked responses was determined by comparing firing rates during the pre-laser baseline and the laser period preceding sound onset across trials (p < 0.05, paired t-test).

Analysis of tone-evoked responses and frequency tuning included only IC neurons with significant excitatory responses to at least one tone frequency and to cortical laser stimulation. For each neuron, the best frequency (BF) was defined as the tone frequency that elicited the highest peak firing rate. To quantify tone-evoked responses, a 15-ms response window was defined around the peak response at the BF (from 5 ms before to 10 ms after the peak). The same response window was then used to quantify tone-evoked firing rates across all tested frequencies.

To quantify cortical modulation of tone-evoked responses, responses were compared between tone-alone and tone-plus-laser trials for each frequency. For both conditions, responses were measured relative to the 20 ms baseline period immediately preceding tone onset. In a subset of transient stimulation cases in which a stable laser baseline could not be obtained, the baseline firing rate was instead estimated from the corresponding time window in separate laser-only trials (n = 10). Frequency tuning curves were constructed from baseline-subtracted tone-evoked firing rates across all tested frequencies. Each tuning curve was smoothed using a three-point Hann window and linearly interpolated by 10-fold. Tuning width was defined as the full width of the tuning curve at half of the maximum firing rate and expressed in octaves.

### Locomotion analysis

Locomotion periods were defined as epochs during which treadmill speed exceeded 2 cm/s. For each IC neuron, significant locomotion-related modulation was determined by comparing firing rates across 1-s segments during stationary and locomotion periods (Student’s t-test, p < 0.05). Mean firing rates during stationary and locomotion periods were then calculated for each neuron. The Modulation Index (MI) for locomotion was defined as (<r>_walking_ - <r>_stationary_)/ (<r>_walking_ + <r>_stationary_), and the MI for laser-evoked responses as (<r>_ON_ - <r>_OFF_)/ (<r>_ON_ + <r>_OFF_), where <r> denotes the mean firing rate of the neuron under indicated condition.

For analysis of modulation timing, locomotion onset was defined as the time point at which the treadmill sensor output changed by 2% of the voltage range (∼3.5 V). Spike times were aligned to locomotion onset, and the average firing rate across locomotion onsets was computed. Neural modulation onset was defined as the first time point within ±0.5 s of locomotion onset at which the smoothed mean firing rate (100 ms hanning window) crossed a threshold set at 2 SD above or below the stationary baseline. The baseline was defined as the period from -1.0 to -0.5 s relative to locomotion onset. For the neurons with a mean baseline firing rate was 0 Hz, a default value of 0.5 Hz was used to estimate the modulation threshold. Modulation latency was calculated as the difference between neural modulation onset and locomotion onset, such that negative values indicate neural modulation preceding locomotion onset. This analysis included only neurons exhibiting clear modulation around locomotion onset and at least 6 locomotion onsets.

For recordings of putative layer 5 and 6 neurons in S1 and M1 during locomotion (Fig. 5j,k), modulation timing analysis was restricted to single units recorded at depths greater than 450 μm in S1 and greater than 400 μm in M1. The depth thresholds corresponded to the depths at which reliable laser-evoked responses from corticocollicular neurons were first observed.

For modulation timing analysis of light-responsive cortical multiunits (Supplementary Fig. 5), spike sorting was performed using template matching in Offline Sorter v4 (Plexon). Direct optogenetic stimulation typically evoked synchronous activity from neighboring cortical neurons, making reliable isolation of individual single units difficult. Laser-evoked waveforms were first inspected to remove stimulation artifacts based on their short latency, characteristic waveform shape, and consistent occurrence at the onset and offset of each laser pulse. For each recording site, the 10–20 largest-amplitude spike waveforms were averaged to generate a template representing the light-responsive multiunit cluster. This template was then applied to both the light-evoked and locomotion recordings from the same recording channel to identify spike waveforms matching the template (fit tolerance = 70 in Offline Sorter).

The mean latency of light-evoked multiunit responses was 2.12 ± 0.18 ms (mean ± SEM), consistent with direct activation of corticocollicular neurons. To verify that the same multiunit cluster was identified under both recording conditions, the similarity between template-matched waveforms from the laser-evoked and locomotion recordings was quantified using Pearson’s correlation coefficient (r = 0.994 ± 0.0019, mean ± SEM). Only multiunit clusters with a signal-to-noise ratio (SNR) > 3 were included in the analysis (SNR = 4.3 ± 0.12, mean ± SEM, n = 12).

### Statistical analysis

Statistical analyses were performed in MATLAB (MathWorks). Normality was assessed using the Shapiro-Wilk test. For normally distributed data, paired or unpaired Student’s t-test were used. For non-normally distributed data, paired comparisons were performed using the Wilcoxon signed-rank test, and unpaired comparisons were performed using the Wilcoxon rank-sum test. Comparisons among multiple groups were performed using the Kruskal–Wallis test. Correlations were assessed using Spearman’s rank correlation coefficient. Statistical significance was set at α = 0.05. Significance levels are denoted as *p < 0.05, **p < 0.01, and ***p < 0.001.

## Data availability

The data that support the findings of this study are available from the corresponding author upon request.

## Acknowledgements

We thank Dr. Won Beom Jung for his assistance in optimizing the MATLAB code for 3D reconstruction.

## Author contributions

Conceptualization, H.Y.K. and G.K.; methodology, H.Y.K., J.H., M.S.-V., and G.K.; investigation, H.Y.K., J.H., and G.K.; writing – original draft, H.Y.K. and G.K.; writing – review and editing, H.Y.K., J.H., and G.K.; supervision, J.L., S.-G.K., and G.K.; funding acquisition, J.L., S.-G.K., and G.K.

## Supplementary figures

**Supplementary Figure 1.**
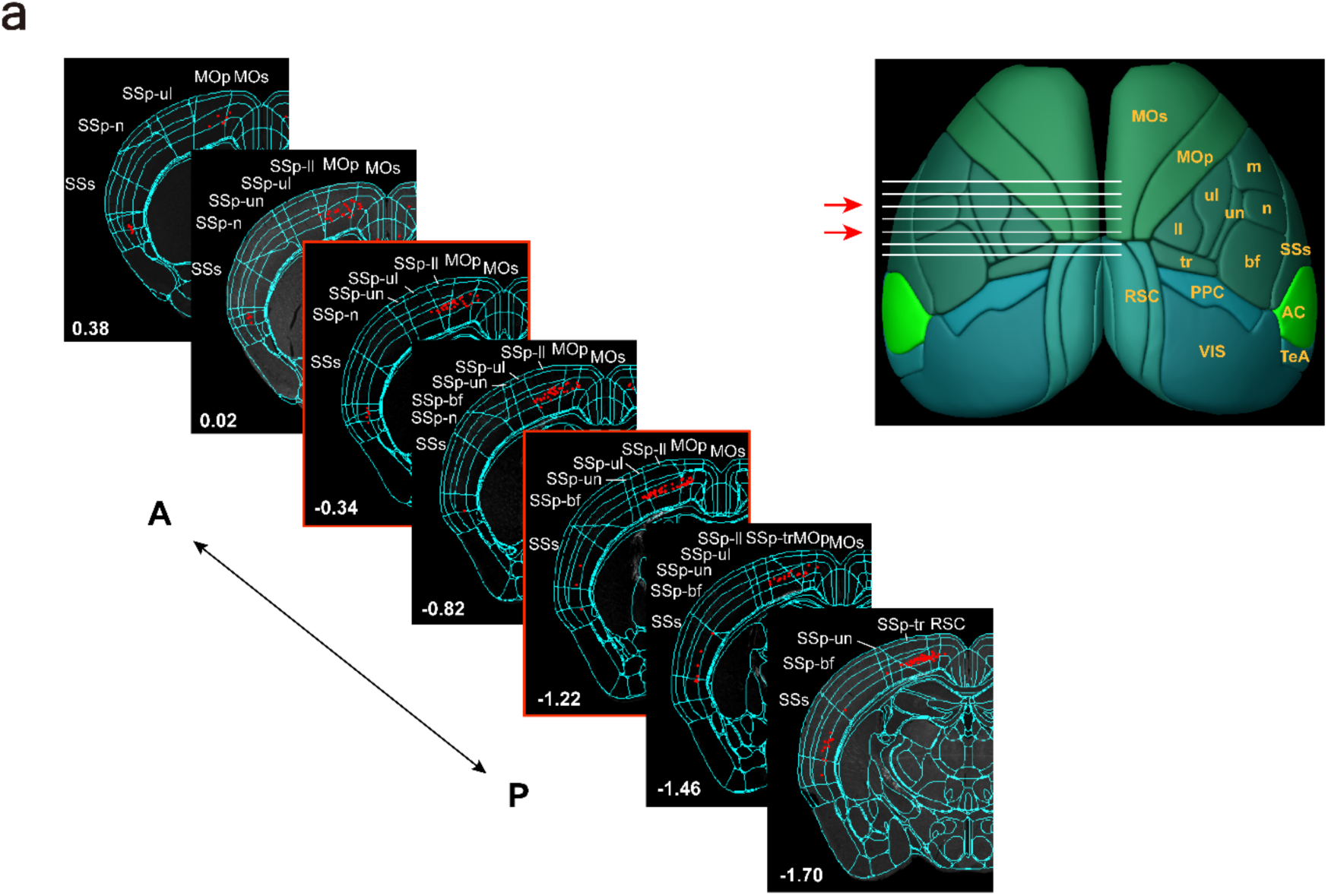
Spatial distribution of IC-projecting neurons in somatosensory and motor cortices. Left: Serial coronal sections arranged along the anterior-posterior (A-P) axis showing reconstructed IC-projecting neurons superimposed on the corresponding Allen Mouse Brain Atlas templates. Highlighted sections (red boxes) correspond to the representative images shown in Fig. 1c. Right: A 3D dorsal view of the mouse cortex showing major cortical regions. White horizontal lines indicate the A-P levels of the coronal sections shown on the left, and red arrows indicate the representative sections highlighted by red boxes. Somatotopic subdivisions of the primary somatosensory cortex (SSp) are labeled according to their body representations: m, mouth; ul, upper limb; ll, lower limb; un, unassigned; n, nose; tr, trunk; and bf, barrel field. MOs: secondary motor cortex; MOp: primary motor cortex; SSp: primary somatosensory cortex; SSs: secondary somatosensory cortex; RSC: retrosplenial cortex; PPC: posterior parietal cortex; AC: auditory cortex; VIS: visual cortex; TeA: temporal association cortex.

**Supplementary Figure 2.**
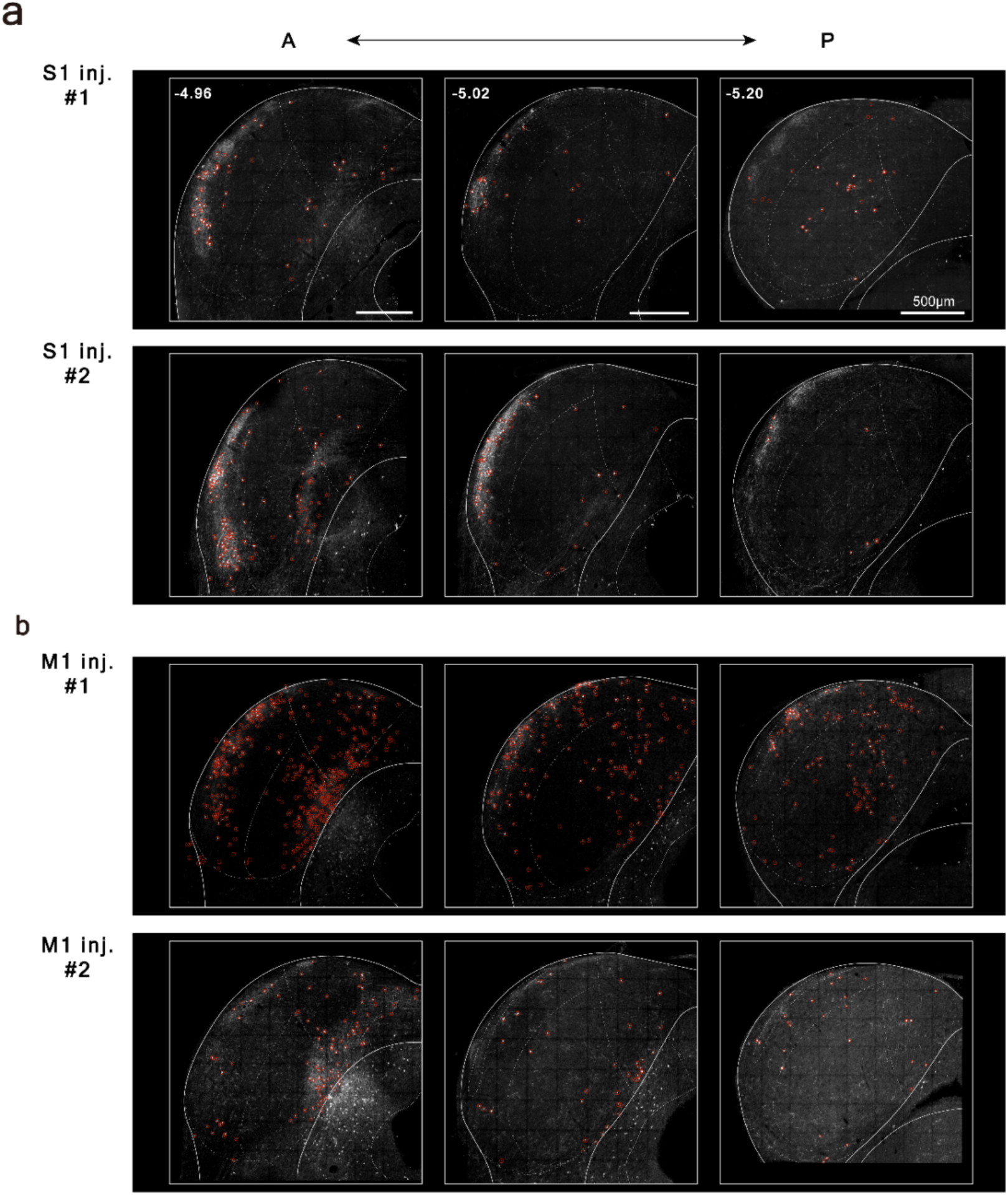
Distribution of transsynaptically labeled recipient IC neurons following S1 and M1 injections. **a-b)** Coronal sections of the IC at 3 rostrocaudal levels following anterograde transsynaptic AAV1-Cre injection into S1 (**a)** or M1 (**b)** of Ai9 reporter mice (n = 2 mice for S1 injection; n = 2 mice for M1 injection). Red circles indicate the manually annotated locations of recipient neurons used for quantification in Fig. 1.

**Supplementary Figure 3.**
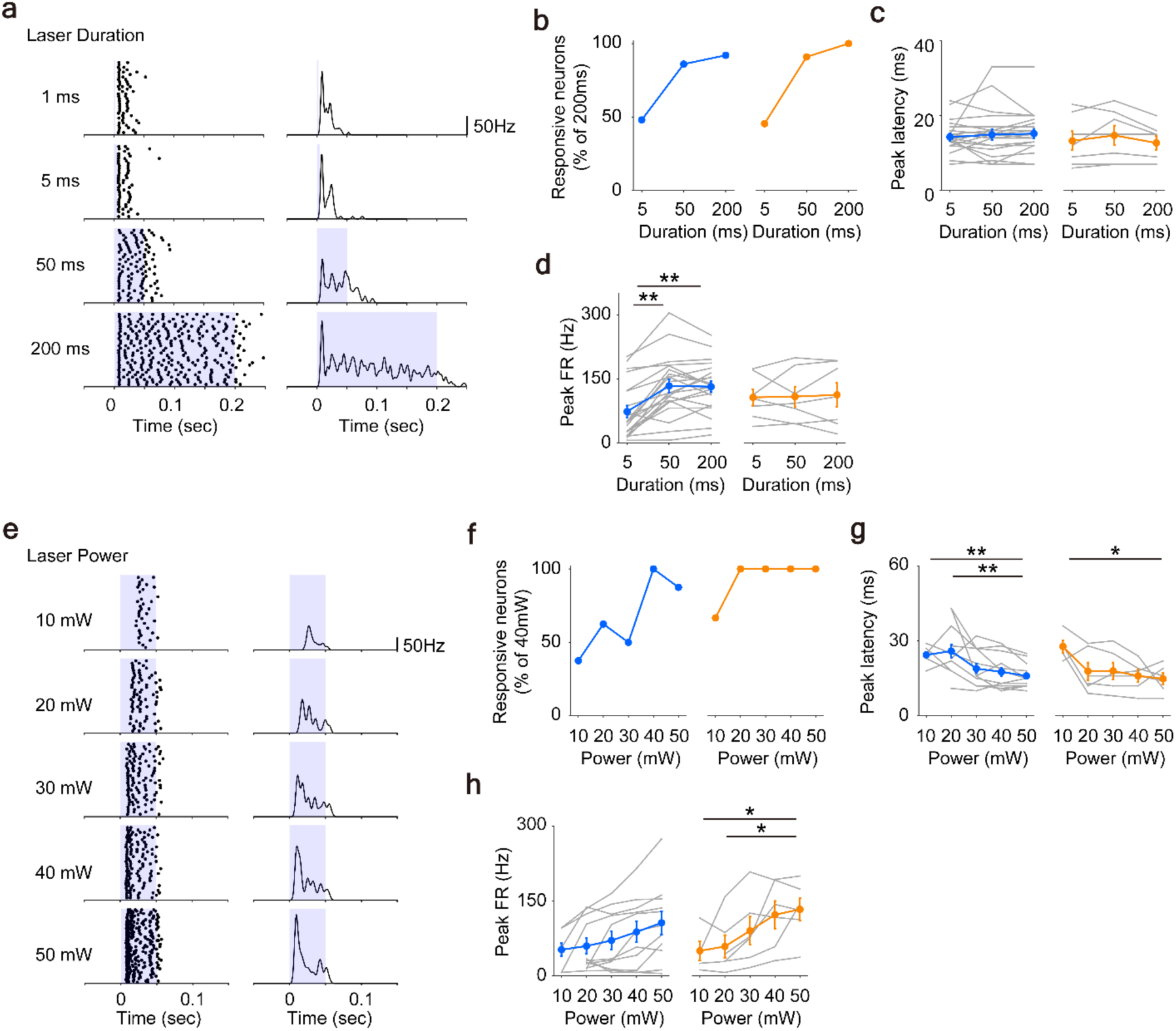
Effects of cortical stimulation parameters on evoked IC activity**. a-d**) Effects of varying laser stimulus duration. **a**) Representative IC responses (raster plots and PSTHs) to M1 stimulation with different laser durations. Blue shading indicates the laser stimulation period. **b**) Proportion of responsive neurons normalized to 200 ms condition (S1, n = 50; M1, n = 22). **c,d**) Peak latency (**c**) and peak firing rate (**d**) as a function of stimulus duration for S1 (blue) and M1 (orange) stimulation (S1, n = 21; M1, n = 9). **e-h**) Effects of varying laser stimulus power. **e**) Representative IC responses to M1 stimulation with different laser powers. Blue shading indicates laser stimulation period. **f**) Proportion of responsive neurons normalized to 40 mW condition (S1, n = 8; M1, n = 6). **g,h**) Peak latency (**g**) and peak firing rate (**h**) as a function of stimulus power for S1 (blue) and M1 (orange) stimulation (S1, n = 12; M1, n = 6). Data are presented as mean ± SEM, with gray lines representing individual neurons. Statistical significance is indicated where applicable (Student’s t-test).

**Supplementary Figure 4.**
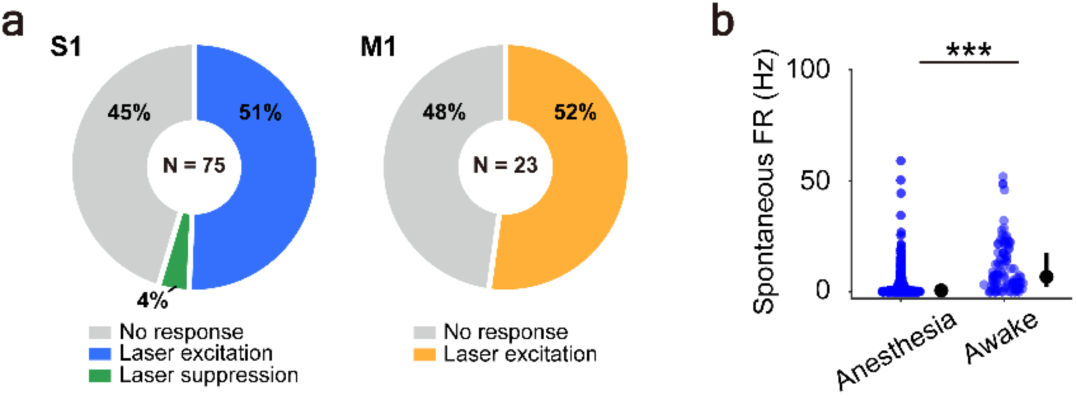
Cortical stimulation responses and spontaneous activity in awake IC recordings. **a)** Proportion of IC neurons exhibiting excitation, suppression, or no response to optogenetic stimulation of S1 (left) or M1 (right) in awake mice. **b)** Comparison of spontaneous firing rates (FR) in anesthetized and awake mice. Median firing rates were 0.67 Hz under anesthesia (n = 340) and 6.81 Hz in awake mice (n = 97; p = 4.51 x 10^-17^, Wilcoxon rank-sum test).

**Supplementary Figure 5.**
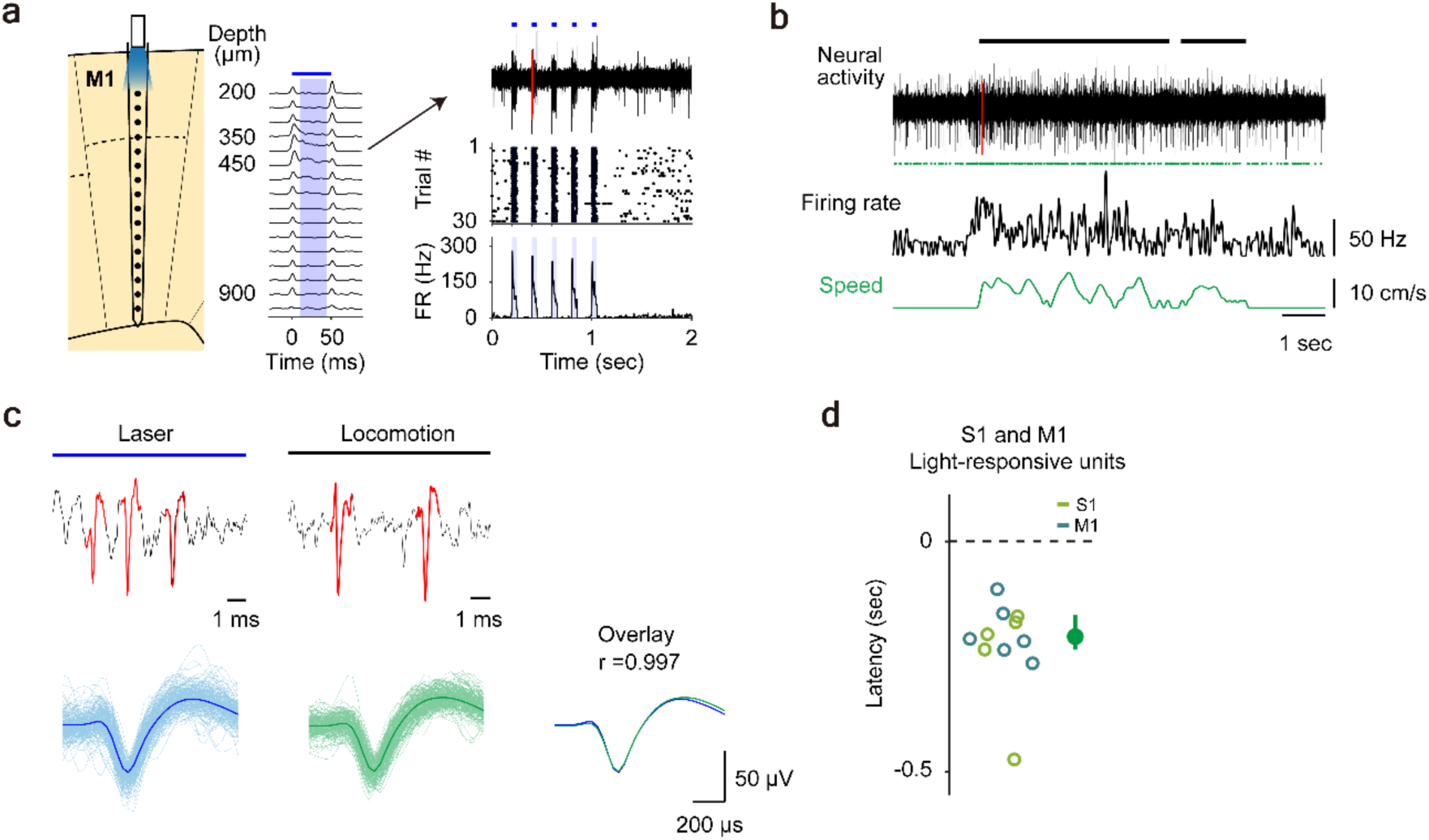
Locomotion-related activity in light-responsive S1 and M1 multiunits. **a**) Representative M1 multiunit recording showing responses to optogenetic stimulation. Left: Schematic of the cortical recording configuration and firing rates recorded at different cortical depths. Blue bar and shading indicate the laser stimulation period. Right: Neural activity, raster plot, and PSTH of the multiunit recorded at 400 μm during laser stimulation. Blue bars above the spike trace indicate laser pulses. **b**) The same multiunit recorded during locomotion. Neural activity, detected multiunit spikes (green dots), smoothed firing rate, and locomotion speed are shown from top to bottom. Black horizontal bars indicate locomotion periods. In (**a**) and (**b**), red-highlighted segments of the neural activity traces are shown at higher magnification in (**c**). **c**) Comparison of laser-evoked and locomotion-related multiunit waveforms (see Methods). Top: Representative waveforms detected during laser stimulation (left) and locomotion (right). Bottom: Superimposed spike waveforms detected during laser stimulation (blue) and locomotion (green), and an overlay of the mean waveforms (right) showing a high degree of similarity between the two conditions (r = 0.997). **d**) Modulation onset latencies relative to locomotion onset for light-responsive S1 and M1 multiunits (median latency = -207 ms; S1: n = 6 units; M1: n = 6 units).

